# H&E-stained Whole Slide Image Deep Learning Predicts SPOP Mutation State in Prostate Cancer

**DOI:** 10.1101/064279

**Authors:** Andrew J. Schaumberg, Mark A. Rubin, Thomas J. Fuchs

**Author notes:** Authors declare no conflicts of interest.

## Abstract

A quantitative model to genetically interpret the histology in whole microscopy slide images is desirable to guide downstream immunohistochemistry, genomics, and precision medicine. We constructed a statistical model that predicts whether or not SPOP is mutated in prostate cancer, given only the digital whole slide after standard hematoxylin and eosin [H&E] staining. Using a TCGA cohort of 177 prostate cancer patients where 20 had mutant SPOP, we trained multiple ensembles of residual networks, accurately distinguishing SPOP mutant from SPOP non-mutant patients (test AUROC=0.74, p=0.0007 Fisher’s Exact Test). We further validated our full metaensemble classifier on an independent test cohort from MSK-IMPACT of 152 patients where 19 had mutant SPOP. Mutants and non-mutants were accurately distinguished despite TCGA slides being frozen sections and MSK-IMPACT slides being formalin-fixed paraffin-embedded sections (AUROC=0.86, p=0.0038). Moreover, we scanned an additional 36 MSK-IMPACT patients having mutant SPOP, trained on this expanded MSK-IMPACT cohort (test AUROC=0.75, p=0.0002), tested on the TCGA cohort (AUROC=0.64, p=0.0306), and again accurately distinguished mutants from non-mutants using the same pipeline. Importantly, our method demonstrates tractable deep learning in this “small data” setting of 20-55 positive examples and quantifies each prediction’s uncertainty with confidence intervals. To our knowledge, this is the first statistical model to predict a genetic mutation in cancer directly from the patient’s digitized H&E-stained whole microscopy slide. Moreover, this is the first time quantitative features learned from patient genetics and histology have been used for content-based image retrieval, finding similar patients for a given patient where the histology appears to share the same genetic driver of disease i.e. SPOP mutation (p=0.0241 Kost’s Method), and finding similar patients for a given patient that does not have have that driver mutation (p=0.0170 Kost’s Method).

**Significance Statement:** This is the first pipeline predicting gene mutation probability in cancer from digitized H&E-stained microscopy slides. To predict whether or not the speckle-type POZ protein [SPOP] gene is mutated in prostate cancer, the pipeline (i) identifies diagnostically salient slide regions, (ii) identifies the salient region having the dominant tumor, and (iii) trains ensembles of binary classifiers that together predict a confidence interval of mutation probability. Through deep learning on small datasets, this enables automated histologic diagnoses based on probabilities of underlying molecular aberrations and finds histologically similar patients by learned genetic-histologic relationships.

Conception, Writing: AJS, TJF. Algorithms, Learning, CBIR: AJS. Analysis: AJS, MAR, TJF. Supervision: MAR, TJF.

---

**G**enetic drivers of cancer morphology, such as E-Cadherin [CDH1] loss promoting lobular rather than ductal phenotypes in breast, are well known. TMPRSS2-ERG fusion in prostate cancer has a number of known morphological traits, including blue-tinged mucin, cribriform pattern, and macronuclei [5]. Computational pathology methods [6] typically predict clinical or genetic features as a function of histological imagery, e.g. whole slide images. Our central hypothesis is that the morphology shown in these whole slide images, having nothing more than standard hematoxylin and eosin [H&E] staining, is a function of the underlying genetic drivers. To test this hypothesis, we gathered a cohort of 499 prostate adenocarcinoma patients from The Cancer Genome Atlas [TCGA]^1^, 177 of which were suitable for analysis, with 20 of those having mutant SPOP (Figs 1, 2, and S1). We then used ensembles of deep convolutional neural networks to accurately predict whether or not SPOP was mutated in the patient, given only the patient’s whole slide image (Figs 3 and 4 panel A), leveraging spatial localization of SPOP mutation evidence in the histology imagery (Fig 4 panels B and C) for statistically significant SPOP mutation prediction accuracy when training on TCGA but testing on the MSK-IMPACT[7] cohort (Fig 5). Further, we scanned 36 additional SPOP mutant MSK-IMPACT slides, training on this expanded MSK-IMPACT cohort and testing on the TCGA cohort. Our classifier’s generalization error bounds (Fig 5 panels A and B), receiver operating characteristic (Fig 5 panels C1 and D1), and independent dataset performance (Fig 5 panels C2 and D2) support our hypothesis, in agreement with earlier work suggesting SPOP mutants are a distinct subtype of prostate cancer [8]. Finally, we applied our metaensemble classifier to the content-based image retrieval [CBIR] task of finding similar patients to a given query patient (Fig 6), according to SPOP morphology features evident in the patient slide dominant tumor morphology.

**Fig. 1.**
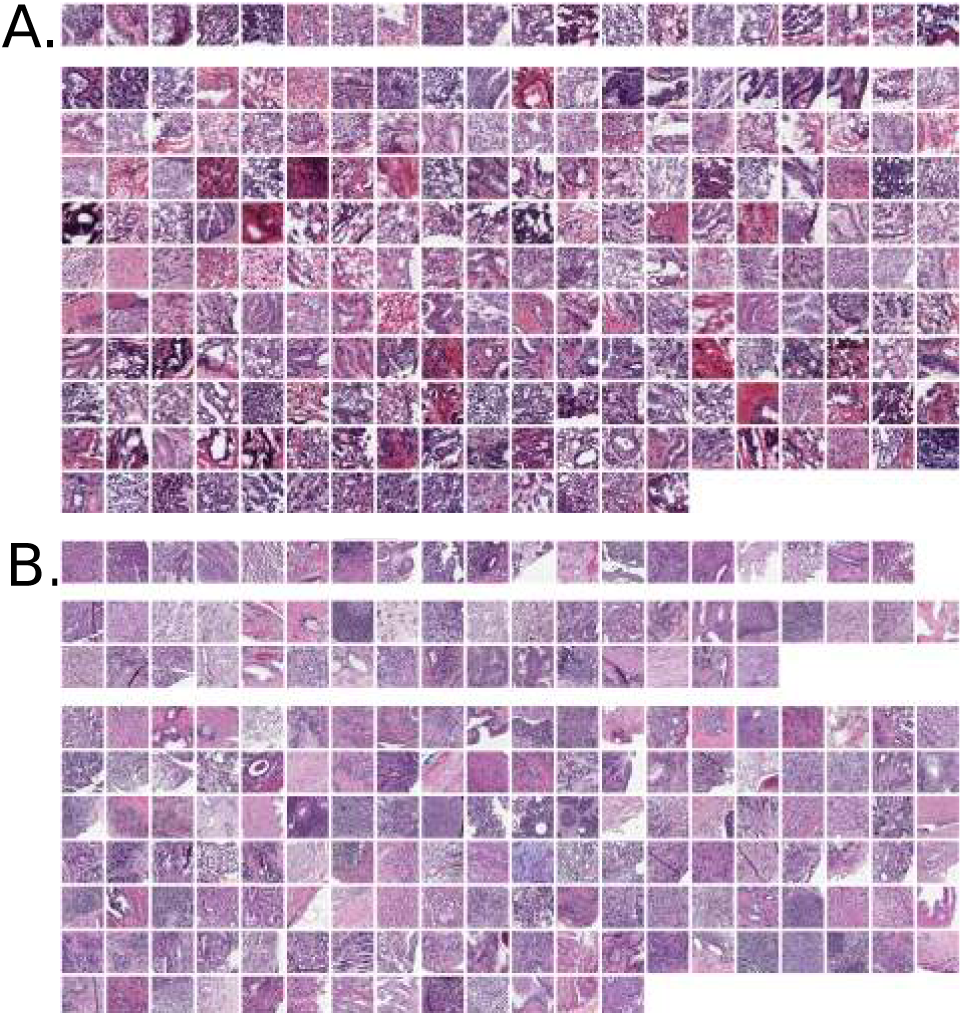
**Panel A:** TCGA cohort of frozen section images. Top row shows 20 SPOP mutants. Bottom rows are 157 SPOP non-mutants, where 25 patients had 2 and 6 patients had 3 acceptable slides available. **Panel B:** MSK-IMPACT cohort of formalin-fixed paraffin-embedded sections, providing higher image quality than frozens. Top row shows 19 SPOP mutants. Middle rows show 36 SPOP mutants scanned as added training data for TCGA testing. Bottom rows are 133 SPOP non-mutants.

**Fig. 2.**
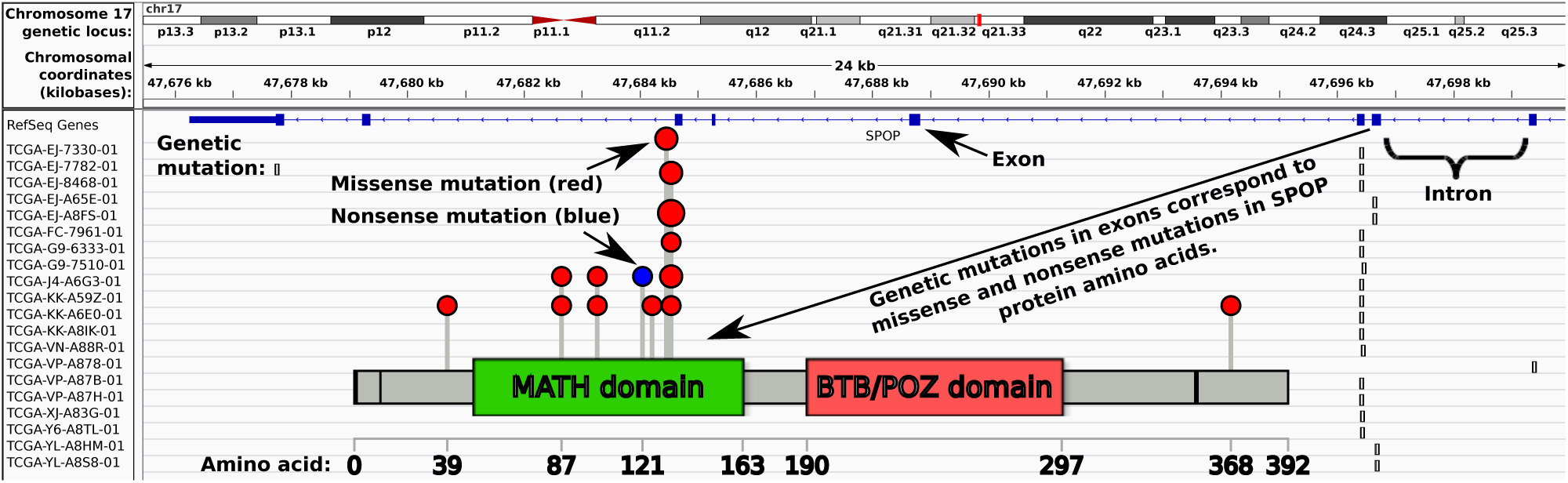
SPOP mutations in the Integrated Genomics Viewer [1, 2], with lollipop plot showing mutations in most frequently mutated domain. For two of twenty patients, somatic SPOP mutations fall outside the MATH domain, responsible for recruiting substrates for ubiquitinylation.

**Fig. 3.**
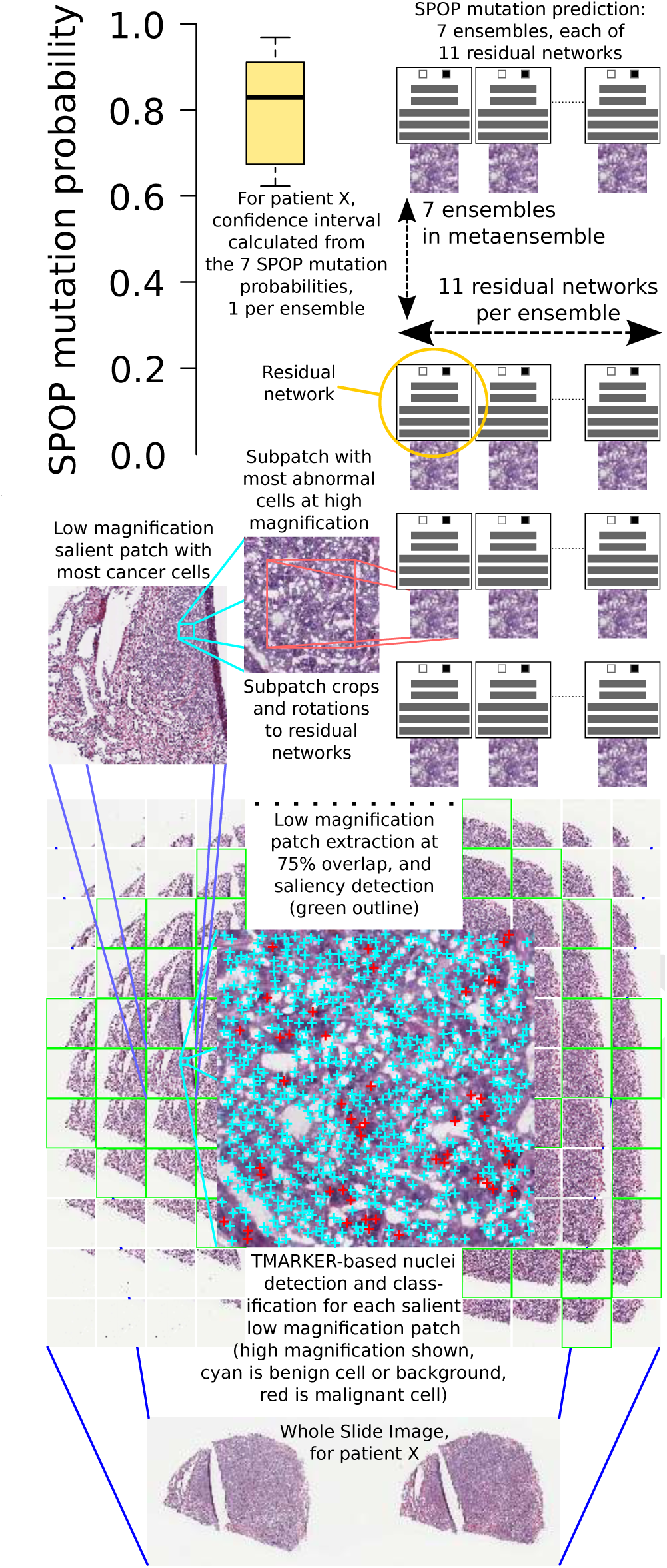
Pipeline: a whole slide image is split into patches (bottom) at low magnification. Salient patches are identified. The salient patch with the most cancer cells is deemed the “dominant tumor” patch and further analyzed. At high magnification, a sliding window within the dominant tumor patch finds the region with maximum abnormal cells. Deep neural networks then predict SPOP mutation and a confidence interval is calculated over these predictions.

**Fig. 4.**
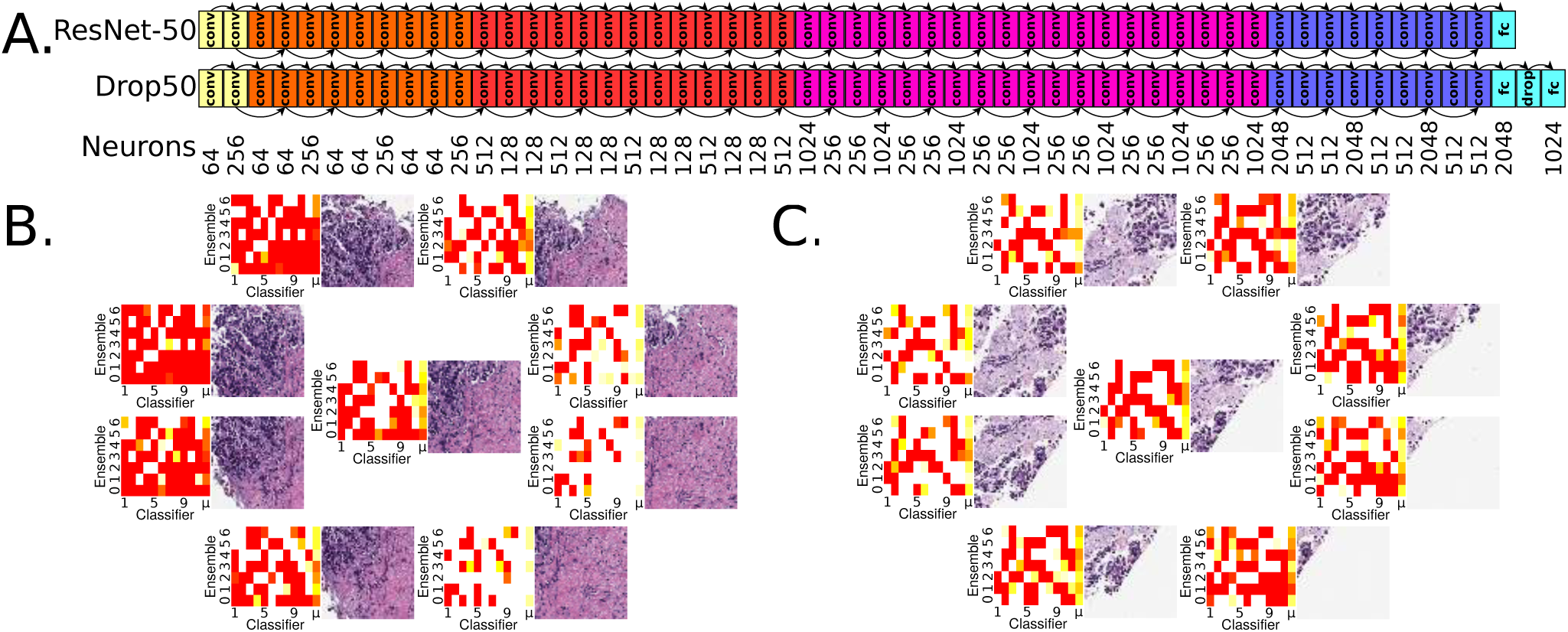
**Panel A:** ResNet-50 architecture [3], top. Customized Drop50 architecture supports ResNet-50 pretraining, but has additional dropout [4] and fully connected neuron layers. In practice at least one of these architectures converges to a validation accuracy of 0.6 or more. Convolutional layers “conv”, fully-connected layers “fc”, 50% dropout layer “drop”. ResNet-50 has 26,560 neurons and Drop50 has 28,574. All seven trials, each trial being a multiarchitectural ensemble of eleven residual networks, is 2,138,630 neurons total for TCGA training and MSK-IMPACT testing, and 2,132,858 neurons total for MSK-IMPACT training and TCGA testing. **Panel B:** Region of interest and surrounding octagon patches, each from an 800×800 pixel [px] patch cropped to 512×512px and scaled to 256×256px. At upper left, histological imagery leads to strong SPOP mutation predictions shown in red. At lower right, no such evidence exists and SPOP mutation is not predicted here, indicated in heatmaps as white rather than red. For each of the nine patches, the weighted mean prediction is calculated, shown in the heatmaps as the *μ* column at right. Each classifier in the ensemble makes a prediction for a patch, and classifiers having greater prediction variance among the nine patches are more weighted in the means for the patches. The metaensemble’s SPOP mutation prediction is 0. 6244, with 95% CI of 0.5218-0.7211 and 99% CI of 0.4949-0.7489, so this patient’s tumor is predicted to have an SPOP mutation at 95% confidence but not 99% confidence. **Panel C:** Another MSK-IMPACT patient shown for the TCGA-trained metaensemble, suggesting there is greater SPOP mutation histological evidence in the lower right. The metaensemble’s SPOP mutation prediction is 0.5528, with 95% CI of 0.5219-0.5867 and 99% CI of 0.5128-0.5962, so there is 99% confidence of SPOP mutation.

**Fig. 5.**
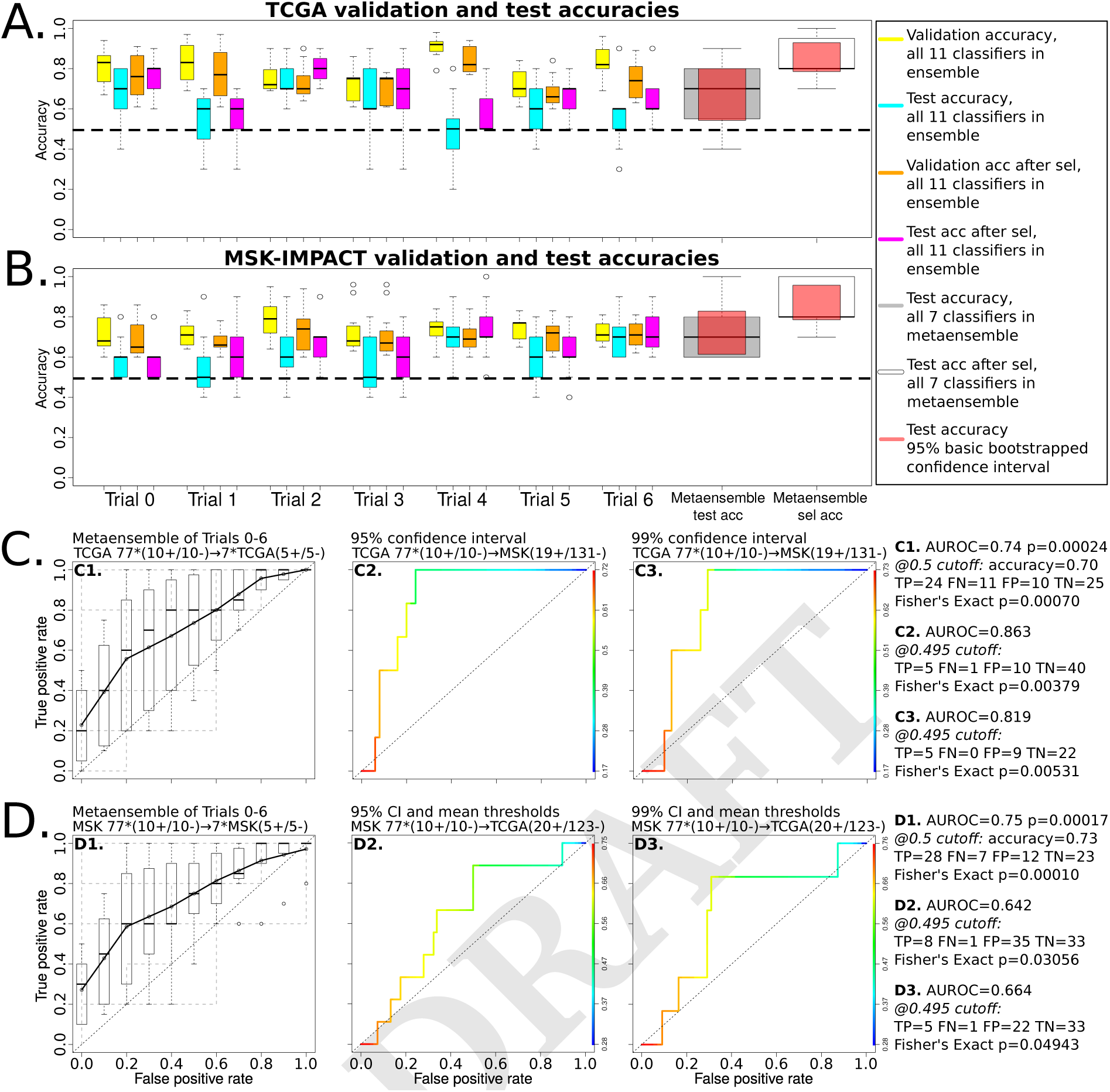
Stricter CIs reduce false negatives and maintain significant Fisher’s Exact p-values, but ignore more predictions as inconclusive (panels C2, C3, D2, D3 – and Alg S3). The metaensemble in panel C1 consists of seven ensembles, with Receiver Operating Characteristics for each shown in Fig S3. The MSK-IMPACT training to TCGA testing performance (0.01 < *p* < 0.05, panels D2 and D3) is expected due to MSK-IMPACT fixed sections being higher quality than TCGA frozen sections (Fig 1). Freezing distorts the slide by effecting watery and fatty tissues differently. Training on distorted data (TCGA) but testing on undistorted data (MSK-IMPACT) appears robust (0.001 < *p* < 0.01, panels C2 and C3) as expected, due to the classifiers learning SPOP-discriminative features despite distortions in the data. This is similar in principle to de-noising autoencoders learning to be robust to noise in their data.

**Fig. 6.**
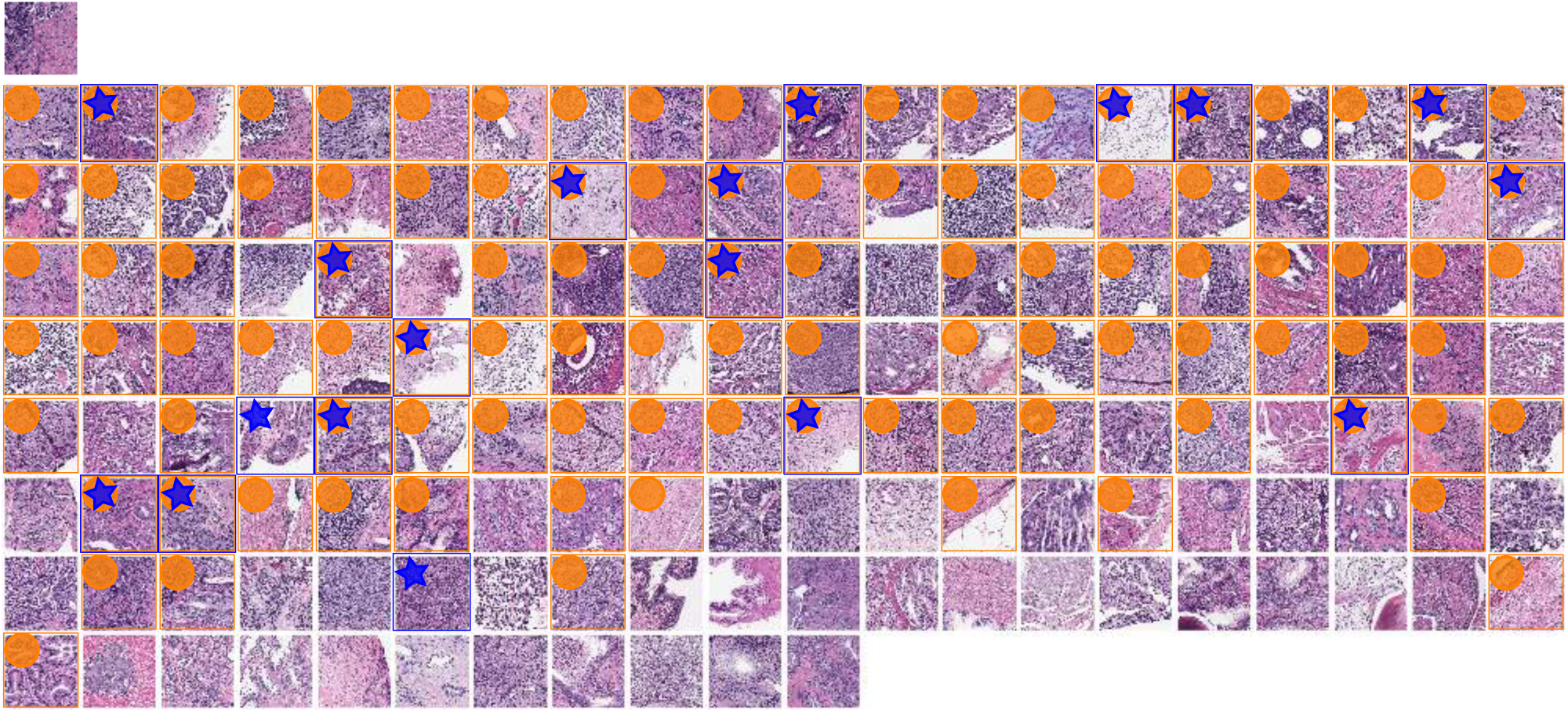
SPOP mutant query image at top left for CBIR using TCGA-trained metaensemble on MSK-IMPACT dataset, showing most similar images in top row, with most similar image leftmost. Less similar images in lower rows, ordered left to right by similarity. One image patch per patient shown. Though the full octagon of patches (Fig 4 panel B, for query) is considered for the query and each retrieved patient, only the center patch is shown here, with a blue surrounding box and star indicating SPOP mutation, while an orange box and circle indicates this retrieved patient’s predicted SPOP mutation 95% CI overlaps with that of the query patient. A 95% CI may in practice usefully limit the number of retrieved results. The most similar patient (top row, leftmost), second most, third, fourth, and fifth have dissimilarity scores of 0.073936, 0.092472, 0.107029, 0.108397, and 0.109131, respectively. The least similar patient (bottom right) has dissimilarity score 0.452150. Mean dissimilarity is 0.219295 with standard deviation 0.075968. Similarity is 1 - dissimilarity. The second most similar patient (top row, second from left) is a mutant, like the query. There are 5 mutants in the top row, 3 in the second, etc, with zero in the bottom row. Patients with SPOP mutation are more similar than patients without SPOP mutation for this query (p=0.02032, one-tailed Mann-Whitney U test).

Previously, pathologists described slide image morphologies, then correlated these to molecular aberrations, e.g. mutations and copy number alterations [9, 10]. Our deep learning approach instead learns features from the images without a pathologist, using one mutation as a class label, and quantifies prediction uncertainty with confidence intervals [CIs] (Fig 3).

Others used support vector machines to predict molecular subtypes in a bag-of-features approach over Gabor filters [11]. The authors avoided deep learning due to limited data available. Gabor filters resemble first layer features in a convolutional network. A main contribution of ours is using pre-training, Monte Carlo cross validation, and heterogeneous ensembles to enable deep learning despite limited data. We believe our method’s prediction of a single mutation is more clinically actionable than predicting broad gene expression subtypes.

Support vector machines, Gaussian mixture modeling, and principal component analyses have predicted PTEN deletion and copy number variation in cancer, but relied on fluorescence *in situ* hybridization [FISH], a very specific stain [12]. Our approach uses standard H&E, a non-specific stain that we believe could be utilized to predict more molecular aberrations than only the SPOP mutation that is our focus here. However, our method does not quantify tumor heterogeneity.

Tumor heterogeneity has been analyzed statistically by other groups [13], as a spatial analysis of cell type entropy cluster counts in H&E-stained whole slides. A high count, when combined with genetic mutations such as TP53, improves patient survival prediction. This count is shown to be independent of underlying genetic characteristics, whereas our method predicts a genetic characteristic, i.e. SPOP mutation, from convolutional features of the imaging.

Clustering patients according to hand-engineered features has been prior practice in histopathology CBIR, with multiple pathologists providing search relevancy annotations to tune the search algorithm [14]. Our approach relies on neither pathologists nor feature engineers, and instead learns discriminative genetic-histologic relationships in the dominant tumor to find similar patients. We also do not require a pathologist to identify the dominant tumor, so our CBIR search is automated on a whole slide basis. Because the entire slide is the query, we do not require human judgement to formulate a search query, so CBIR search results may be precalculated and stored for fast lookup.

## Results

### Molecular information as labels of pathology images opens a new field of molecular pathology

Rather than correlating or combining genetic and histologic data, we predict a gene mutation directly from a whole slide image with unbiased H&E stain. Our methods enable systematic investigation of other genotype and phenotype relationships, and serve as a new supervised learning paradigm for clinically actionable molecular targets, independent of clinician-supplied labels of the histology. Epigenetic, copy number alteration, gene expression, and post-translational modification data may all label histology images for supervised learning. Future work may refine these predictions to single-cell resolution, combined with single-cell sequencing, immunohistochemistry, FISH, mass cytometry[15], or other technologies to label corresponding H&E images or regions therein. We suggest focusing on labels that are clinically actionable, such as gains or losses of function.

### SPOP mutation state prediction is learnable from a small set of whole slides stained with hematoxylin and eosin

Despite SPOP being one of the most frequently mutated genes in prostate adenocarcinomas[8], from a TCGA cohort of 499 patients only 177 passed our quality control (Alg S1) and only 20 of these had SPOP mutation. Meanwhile in the 152-patient MSK-IMPACT cohort there were only 19 SPOP mutants, and though we could scan an additional 36 SPOP mutant archived slides, there are difficulties in practice acquiring large cohorts of patients with both quality whole slide cancer pathology images and genetic sequencing represented. This challenge increases for rare genetic variants. Moreover, different cohorts may have different slide preparations and appearances, such as TCGA being frozen sections and MSK-IMPACT being higher quality formalin-fixed paraffin embedded sections (Fig 1). Nonetheless, our pipeline (Fig 3) accurately predicts whether or not SPOP is mutated in the MSK-IMPACT cohort when trained on the TCGA cohort (Fig 5), and vice versa. We leverage pretrained neural networks, Monte Carlo cross validation, class-balanced stratified sampling, and architecturally heterogeneous ensembles for deep learning in this “small data” setting of only 20 positive examples, which may remain relevant for rare variants as more patient histologies and sequences become available in the future.

### SPOP mutation state prediction is accurate and the prediction uncertainty is bounded within a confidence interval

Our pipeline’s accuracy in SPOP mutation state prediction is statistically significant: (i) within held-out test datasets in TCGA,

(ii) within held-out test datasets in MSK-IMPACT, (iii) from TCGA against the independent MSK-IMPACT cohort (Fig 5), and (iv) vice versa. The confidence interval [CI] calculated using seven ResNet ensembles allows every prediction to be evaluated statistically: is there significant evidence for SPOP mutation in the patient, is there significant evidence for SPOP non-mutation in the patient, or is the patient’s SPOP mutation state inconclusive. Clinicians may choose to follow the mutation prediction only if the histological imagery provides acceptably low uncertainty.

### SPOP mutation state prediction is automated and does not rely on human interpretation

Unlike Gleason score, which relies on a clinician’s interpretation of the histology, our pipeline is automated (Fig 3) – though detecting overstain, blur, and mostly background in the slide are not yet automated (Alg S1). The pipeline’s input is the whole digital slide and the output is the SPOP mutation prediction bounded within 95% and 99% CIs. Moreover, our pipeline does not require a human to identify a representative region in the slide, as is done to create tissue microarrays [TMAs] from slides.

### SPOP mutation state prediction finds patients that appear to share the same genetic driver of disease

Each ensemble (Fig 3) may be used as a feature to predict patient similarity, where similar patients on average share the same ensemble-predicted SPOP mutation probability (Fig 6). A patient’s histology may be studied in the context of similar patients’ for diagnostic, prognostic, and theragnostic considerations.

### Molecular pathology, such as characterizing histology in terms of SPOP mutation state, leads directly to precision medicine

For instance, non-mutant SPOP ubiquitinylates androgen receptor [AR], to mark AR for degradation, but mutant SPOP does not. Antiandrogen drugs, such as flutamide and enzalutamide, promote degradation of AR to treat the cancer, though mutant AR confers resistance [16, 17].

### SPOP mutation state prediction provides information regarding other molecular states

SPOP mutation is mutually exclusive with TMPRSS2-ERG gene fusion [8], so our SPOP mutation predictor provides indirect information regarding the TMPRSS2-ERG state and potentially others.

## Discussion

We summarize a whole slide with a maximally abnormal subregion within the dominant tumor, such that the vast majority of the slide is not used for deep learning. Tissue microarrays take a similar though manual approach, using a small circle of tissue to represent a patient’s cancer. In a sense, our patch extraction, saliency prediction, and TMARKER-based cell counting pipeline stages together model how a representative patch may be selected to identify the dominant cancer subtype in the slide overall, ignoring other regions that may not have the same genetic drivers. This subregion has many desirable properties: (i) it is salient at low magnification, i.e. diagnostically relevant to a pathologist at the microscope [18], (ii) it has the maximum number of malignant cells at low magnification, which is a heuristic to locate the dominant tumor, presumably enriched for cells with driver mutations such as SPOP due to conferred growth advantages and reflected in the bulk genetic sequencing we use as mutation ground truth, (iii) it has the maximum number of abnormal cells at high magnification, which is a heuristic to locate a subregion with most nuanced cellular appearance, presumably concentrated with cellular-level visual features that can discriminate between cancers driven by SPOP versus other mutations, and (iv) at 800×800 pixels, it is large enough at high magnification to capture histology, e.g. gland structure.

A holistic approach that considers for deep learning patches spatially distributed widely throughout the slide, rather than only the dominant tumor, could improve performance and provide further insight. Gleason grading involves such a holistic approach, identifying first and second most prevalent cancer morphologies in a whole slide. The complexities of multiple and varying counts of representatives per slide are future work.

We use multiple classifier ensembles to predict the SPOP mutation state. Though each residual network [3] classifier tends to predict SPOP mutation probability in a bimodal distribution, i.e. being either close to 0 or close to 1, averaging these classifiers within an ensemble provides a uniform distribution representing the SPOP mutation probability (Fig S2).

Deep learning typically requires large sets of training data, yet we have only 20 patients with SPOP mutation, nearly an order of magnitude fewer than the 157 patients without somatic SPOP mutation in the TCGA cohort. Deep learning in small data is a challenging setting. We confront this challenge by training many residual networks [ResNets] on small draws of the data (Alg S2), in equal proportions of SPOP mutants and SPOP non-mutants, then combining the ResNets as weak learners, to produce a strong learner ensemble [19, 20], similar in principle to a random forest ensemble of decision trees [21]. Empirically, the great depth of ResNets appears to be important because CaffeNet [22] – a shallower 8-layer neural network based on AlexNet [23] – so rarely achieved validation accuracy of 0.6 or more predicting SPOP mutation state than an ensemble could not be formed.

Pretraining the deep networks is essential in small data regimes such as ours, and we use the ResNet-50 model pretrained on ImageNet [3]. We used both the published ResNet-50 and our customized ResNet-50 that included an additional 50% dropout [4] layer and a 1024-neuron fully-connected layer. In practice, for difficult training set draws, often at least one of these architectures converged to validation accuracy of 0.6 or more. For data augmentation, the 800×800px images at high magnification were trimmed to the centermost 512×512px in 6 degree rotations, scaled to 256×256px, flipped and unflipped, then randomly cropped to 224×224 within Caffe [22] for training. Learning rates varied from 0.001 to 0.007, and momentums from 0.3 to 0.99. Test error tended to be worse with momentum less than 0.8, though low momentum allowed convergence for difficult training sets. Learning rates of 0.001, 0.0015, or 0.002 with momentum of 0.9 was typical. We used nVidia Titan-X GPUs with 12GB RAM for training. At 12GB, one GPU could train either ResNet architecture using a minibatch size of 32 images.

## Materials and Methods

Please see Section S1 for discussion of (i) digital slide acquisition and quality control, (ii) region of interest identification, (iii) data augmentation, (iv) deep learning, and (v) cross validation.

### Ensemble aggregation

To estimate generalization error, the 11 classifiers in a trial were selected by highest validation accuracy to form an ensemble, and a metaensemble from 7 independent trial ensembles. A 95% basic bootstrapped CI^2^ indicated generalization accuracy from these 7 trials was 0.58-0.86 on TCGA (Fig 5 panel A) and 0.61-0.83 on MSK-IMPACT (Fig 5 panel B), – both significantly better than 0.5 chance. On TCGA, metaensemble AUROC was 0.74 (p=0.00024), accuracy was 0.70, and Fisher’s Exact Test p=0.00070 (Fig 5 panel C1). On MSK-IMPACT, metaensemble AUROC was 0.75 (p=0.00017), accuracy was 0.73, and Fisher’s Exact Test p=0.00010 (Fig 5 panel D1). These two single-dataset estimates indicate the method performs significantly better than chance on unseen data. We formed tuned ensembles (Fig S4), where 11 classifiers in a trial were selected by highest ensemble *test* accuracy, which no longer estimates generalization error but promotes correct classification on average across ensembled classifiers for each test example (Fig 5, panels A and B, white versus gray at right).

### Independent evaluation

We tested the TCGA-trained metaensemble of 7 tuned 11-classifier ensembles on the MSK-IMPACT cohort, and vice-versa. Each patient had one slide. For each slide, a central region of interest and surrounding octagon of patches were found, for nine patches per slide (Fig 4 panels B and C). We hypothesized that SPOP mutation leads to 1-2 nine-patch-localized lesions, meaning adjacent patches have similar classifier-predicted SPOP mutation probability. Therefore, we first calculated the SPOP mutation prediction weighted mean over all classifiers in an ensemble, with classifiers having greater nine-patch prediction variance being more weighted in the mean. Second, of the nine patches we formed all possible groups of three adjacent patches and assigned to a group the minimum SPOP mutation prediction weighted mean of any patch in the grouped three, then took the second-greatest group as a classifier’s overall patient cancer SPOP mutation prediction. Within an ensemble, we weighted a classifier’s overall prediction by the classifier’s nine-patch mutation prediction variance, because classifiers with high variance (e.g. mostly mutant, but 1-3 of nine non-mutant) reflect the lesion hypothesis better than classifiers that predict all non-mutant (0) or all mutant (1). Thus on MSK-IMPACT, the TCGA metaensemble assigned stochastically greater scores to mutants than non-mutants (AU-ROC 0.86) and accurately distinguished mutants from non-mutants (Fisher’s Exact p=0.00379) (Fig 5 panel C2). Moreover on TCGA, the MSK-IMPACT metaensemble assigned stochastically greater scores to mutants than non-mutants (AUROC 0.64) and accurately distinguished mutants from non-mutants (Fisher’s Exact p=0.03056) (Fig 5 panel D2).

### CBIR evaluation

The metaensemble consists of seven ensembles. Each ensemble predicts SPOP mutation probability as a uniformly distributed random variable (Fig S2). Each ensemble is tuned to a different test set (Sec S2 and Table S1), so we treat each ensemble’s prediction as a feature, for seven 32-bit SPOP CBIR features total. For CBIR, a patient’s dissimilarity to the query patient is the mean of absolute differences in these seven features, e.g. a dissimilarity of 0.2 means on average an ensemble predicts the patient’s SPOP mutation probability is 0.2 different than the query. A similarity score is 1 minus dissimilarity. We evaluated the TCGA-trained metaensemble on each of the 19 SPOP mutants in MSK-IMPACT (Fig 1), for each mutant calculating a CBIR p-value, with a low p-value indicating it is not due to chance alone that dissimilarities of mutants were lower than non-mutant dissimilarities (Fig 6). To conduct a metaanalysis of these 19 dependent p-values, we may use neither (a) Fisher’s Method [24] because our p-values are not independent, nor (b) Empirical Brown’s Method [25] because 100 p-values are required for convergence, so we used Kost’s Method^3^ [26]. Kost’s Method found an overall p=0.024076 (14.6 degrees of freedom [d.f.]), so it is not due to chance alone that our CBIR returns SPOP mutant patients with lower dissimilarity scores than non-mutants for each SPOP mutant patient query. Similarly, our CBIR tool returned non-mutant patients with lower dissimilarity than SPOP mutants for each of the 133 MSK-IMPACT non-mutant patients (p=0.016960 Kost’s Method, 22.2 d.f.).

## Acknowledgments

AJS was supported by NIH/NCI grant F31CA214029 and the Tri-Institutional Training Program in Computational Biology and Medicine (via NIH training grant T32GM083937). TJF was supported by an MSK Functional Genomics Initiative grant. This research was funded in part through the NIH/NCI Cancer Center Support Grant P30CA008748. We gratefully acknowledge NVIDIA corporation for GPU support under the GPU Research Center award to TJF. TJF is a founder, equity owner, and Chief Scientific Officer of Paige.AI.

## Supporting Information

### S1. Supplementary materials and methods

#### Digital slide acquisition and quality control

TCGA slides and SPOP mutation state was downloaded from cBioPortal [27]. The MSK-IMPACT cohort was available internally. We discarded slide regions that were not diagnostically salient [18]. We discarded slide regions that were (i) over-stained eosin blue, (ii) > 50% erythrocytes, (iii) > 50% empty, or (iv) > 50% blurred, by inspection (Alg S1). Patients having no regions remaining were discarded. A region is a 800×800 pixel [px] patch at 8 microns per pixel *[μpp]*, ~100x magnification.

#### Region of interest identification

For each region, we used TMARKER[28] to identify cell nuclei and classify them as benign, malignant, or unknown – using a corresponding 12750×12750px patch at level 0 of the slide SVS file (0.25 *μpp*) for the 800×800px region passing quality control (Fig 3). The 12750×12750px patch having the most malignant cells in this slide was deemed the “dominant tumor” after searching in a 10px grid for an 800×800px region of interest patch (at 4 *μpp*) having maximal abnormal cells. A malignant cell is 1 abnormal cell while an unknown cell is 0.5 abnormal cell and 0.5 normal cell. We expected a mutant driver gene, such as SPOP, to confer a growth advantage evident in the region of interest and show histological features, such as ducts. By maximizing abnormal cell counts, we intended to find as many discriminative image textures based on SPOP-mutation-driven cancer cells as possible.

#### Data augmentation

In an 800×800px region of interest patch, the centermost 512×512px patch was selected, rotated in six degree increments for rotational invariance, and scaled to 256×256px. Eight 800×800px patches were also selected in a surrounding octagon formation around the region of interest, offset 115 and 200px away from the central region of interest, so two adjacent octagon patches were 230px away from each other and 230px away from the central patch (Fig 4 panels B and C). These patches were rotated and scaled. This integrates a circular area to summarize the slide, akin to circular tissue cuts for TMA spots.

#### Deep learning

We used Caffe [22] for deep learning a binary classification model given the 256×256px patches labeled by SPOP mutation state. We adapted the ResNet-50 [3] model pre-trained on ImageNet [29] data by re-initializing the last layer’s weights (Fig 4 panel A), then trained on pathology images. Classifier predictions followed a sharply bimodal distribution of 0 or 1, but ensemble predictions followed a uniform distribution suitable for CIs (Fig S2).

#### Cross validation

For a trial, a test set of 5 positive and 5 negative [5+/5-] examples was used, where a positive example is a patient with SPOP mutation (Table S1). If a patient had multiple slides that passed quality control, at most one was used in the trial, taken at random. With the test set held out, we ran Monte Carlo cross validation 13 times, each time (i) drawing without replacement a training set of 10+/10- examples and validation set of 5+/5- examples, and (ii) training both a ResNet-50 and Drop50 classifier (Fig 4 panel A). Our convergence criterion was classifier validation accuracy >= 0.6 with 10000 – 100000 training iterations. An ensemble was 11 classifiers, one per Monte Carlo cross validation, allowing 2 of 13 Monte Carlo cross validation runs to converge poorly or not at all (Alg S2). Empirically, this class-balanced stratified-sampling ensemble approach reduces classifier bias towards the majority class and broadly samples both classes.

##### Algorithm S1

Preprocessing: Each patient’s SPOP mutation state is paired with the whole slide image patch having the maximum number of abnormal cells, where abnormal is either cancer or unknown cell type, rather than healthy cell type. This patch is taken from the dominant tumor patch. The dominant tumor patch is both salient and has the maximum number of cancer cells. The dominant tumor patch is at low magnification, while the abnormal patch is at high magnification within the dominant.

for all *prostate adenocarcinoma patients* do *spop* → *SPOP normal/mutated state as* 0/1 *slide* → *whole slide image of cancer biopsy patches* → 75% *overlap 800×800px images* → *slide salient patches* → *predict_saliency*(*patches*) *pri_tumor_patch* → *max_cancer* (*salient_patches*) *abn_patch* → *max_abnormal*(*pri_tumor_patch*) if *abn_patch* < 50% *blurred* then if *abn_patch* < 50% *background* then if *abn_patch* < 50% *erythrocytes* then if *abn_patch not eosin overstained* then *append* (*abn_patch, spop*) *to data_set* return *data_set*

##### Algorithm S2

Training: Train a residual network on the abnormal patch representing each patient, labeled with SPOP mutation state. The final predictor is an ensemble of eleven of such ResNets. After drawing a test set, training and validation sets are drawn from the remaining patients in the preprocessed data set (Alg S1). Within a Monte Carlo cross validation run, training and validation sets do not overlap. All draws are without replacement. Thirteen Monte Carlo cross validation runs are attempted in parallel, where training stops after eleven have validation accuracy of 0.6 or more. Some runs do not achieve 0.6 after many attempts. See Fig 4 for architecture details of r50 as “ResNet-50” and *drp5* as “Drop50”.

for *trial in* 0,1, 2, 3, 4 do *test_set* → *draw*(5 *SPOP mutants*, 5 *SPOP normal*) for *monte in* 1, 2,…,13 *in parallel* do *train_set* → *draw*(10 *mutants*, 10 *normal*) *valid_set* → *draw*(5 *mutants*, 5 *normal*) *testable_model* → *null* repeat *r*50_*model* → *train_r*50(*train_set,valid_set*) *drp5_model* → *train_drp5*(*train_set,valid_set*) *r*50_*acc* → *accuracy* (*valid_set*, *r*50_*model*) *drp*5_*acc* → *accuracy* (*valid_set*, *drp*5_*model*) if *r*50_*acc* >= 0.6 then *testable_model* → *r*50_*model* else if *drp*5_*acc* >= 0.6 then *testable_model* → *drp*5_*model* until 11 *testable models* or *testable_model* = *null append testable_model to testable_models ensemble* → *testable_models with ensemble averaging ensemble_acc* → accuracy (*test_set, ensemble*) *append ensemble_acc to ensemble_accs append ensemble to ensembles* return (*ensembles, ensemble_accs*)

##### Algorithm S3

Prediction: Use each of 7 ensembles to predict SPOP mutation state, then compute a bootstrapped CI of SPOP mutation state given these predictions. Each prediction is a probability of SPOP mutation in the patient. If this probability is sufficiently high to surpass a defined threshold, the patient’s cancer is predicted to have an SPOP mutation. Likewise, if this probability is sufficiently low to fall below a defined threshold, the patient’s cancer is predicted to not have an SPOP mutation relative patient germline. Probabilities between these upper and lower thresholds are inconclusive predictions. Threshold calibration depends both on the dataset and the cost of false positives/negatives. For instance, if the CI lower bound is > 0.5, there is significant confidence that the patient has SPOP mutation. If the CI upper bound is < 0.5, there is significant confidence that the patient does not have SPOP mutation. Otherwise, the patient’s SPOP mutation state cannot be confidently predicted. When training on TCGA and testing on MSK-IMPACT with a 99% CI (Fig 5 panel C3), the metaensemble did not commit any false negatives at a 0.5 cutoff, so we calibrated the cutoff to 0.495 to make one more true positive while still not committing false negatives. Thus SPOP mutation was predicted when the lower CI bound was above 0.495, and SPOP was predicted to not be mutated when the upper CI bound was below 0.495, with the remaining cases being inconclusive (Fig 5 panels C2 and C3). When training on MSK-IMPACT and testing on TCGA, false positives were an issue, presumably because TCGA frozen sections are poorer image quality than MSK-IMPACT fixed sections (Fig 1), so we changed the lower threshold to the mean, converting many false positives to true negatives. Thus SPOP mutation was predicted when the lower CI bound was above 0.495, and SPOP was predicted to not be mutated when the metaensemble mean was below 0.495 (Fig 5 panels D2 and D3), with the remaining cases being inconclusive. In this way, the CI bounds or mean served as thresholds in a calibrated dataset-dependent manner, and we recommend such calibration when testing on a new dataset. Section S3 defines *P*_*indep*_(*X*|*I*_0_, *I*_1_, …, *I*_8_, *C*_1_, *C*_2_, …, *C*_11_), which calculates an overall SPOP mutation prediction for the patient given the central patch, surrounding octagon of patches (Fig 4 panels B and C), and 11 classifiers in an ensemble.

*slide* → *whole slide image of cancer biopsy patches* → 75% *overlap* 800*x*800*px images* → *slide salient patches* → *predict_saliency*(*patches*) *pri_tumor_patch* → *max_cancer* (*salient_patches*) *abn_patch* → *max_abnormal*(*pri_tumor_patch*) for *ensemble in ensembles* do *I*_0_, *I*_1_, …, *I*_8_ → *abn_patch and surrounding octagon spop_prediction* → *P*_*indep*_(*X*|*I*_0_, *I*_1_,…, *I*_8_, *C*_1_, *C*_2_, …,*C*_11_) *append spop_prediction to spop_predictions* return *confidence_interval*(*spop_predictions*)

### S2. TCGA dataset learning

See Fig 5 panel C1 for TCGA testing of the TCGA-trained metaensemble, with Fig S4 showing TCGA-trained tuned metaensemble performance. Training was as follows:

1. *P*_*learn*_(*X* = *Mutant*) = *P*_*learn*_(*X* = *Normal*) = 0.5. Each classifier is trained on twenty patients, with ten having mutant SPOP with respect to the patient germline. With this balanced dataset for training, the probability that a training example is a mutant is the same as the probability that a training example is normal, 0.5. This differs from the proportion of patients having a somatic SPOP mutation, ~10% in the TCGA dataset. Classifiers that learn on unbalanced datasets may simply learn to predict the predominant class.
2. Learning *P*_*learn*_(*X*|*I*, *C*_1_, *C*_2_,…, *C*_11_) for each patient representative histology image I and eleven classifiers C. This learns the probability that a classifier will predict the patient is mutated, given the representative histology image of the patient. An ensemble of an infinite number of classifiers would give the expected value of this probability. We use eleven classifiers. Each classifier tends to emit either a zero or one, predicting mutant or normal, i.e. *P*_*learn*_(*X*|*I*, *C*) tends to be 0 or 1. We take the ensemble average to be the probability *P*_*learn*_, which tends to be uniformly distributed rather than bimodal (Fig S2): 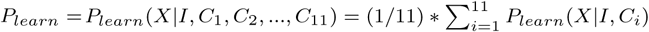.
3. Training and validation sets consist of a central image patch and eight surrounding image patches in an octagon formation, where the distance between the central patch and a surrounding patch is the same as the distance between any two adjacent surrounding patches. All patches had the same label, either 1 for patients with somatic SPOP mutation or 0 for patients without somatic SPOP mutation. This approximately circular region of nine patches per patient provides more information for training, more closely resembles the region available from a circle of tissue in a TMA, and does not sample far away tissues that may have different molecular drivers of disease.
4. The test set consists of a single patch per patient, without a surrounding octagon of eight patches. Test accuracy and squared loss were optimized from the converged trained models having at least 0.6 validation accuracy, i.e. the “sel” ensembles in Fig S4. This optimization selected at most one model from 11 of 13 Monte Carlo cross validation runs, where the selection of 11 models for the ensemble had the highest accuracy and lowest squared loss on the test set. This (a) ensures on unseen data the ensemble has both the discriminative power and diversity to correctly predict *P*_*learn*_, which is a function of a single image, and (b) optimizes each ensemble according to a different test set, with the aim of decorrelating ensembles for a more general metaensemble, much like how decision trees should be decorrelated for a more general random forest [21].

### S3. MSK-IMPACT dataset testing

See Fig 5 panels C2 and C3 for MSK-IMPACT testing of the TCGA-trained metaensemble. Testing was as follows, and the analogous procedure holds for TCGA testing of the MSK-IMPACT-trained metaensemble (Fig 5 panels D2 and D3):

1. 9* *P*_*indep*_(*X* = *Mutant*) ≈ *P*_*indep*_(*X* = *Normal*). The ensembles learned on the TCGA dataset are tested on an independent MSK-IMPACT dataset 152 patients, 19 having somatic SPOP mutation, so the probability that one of these MSK-IMPACTed patients is mutated is only ~10%. This is in sharp contrast to the TCGA training set, where the probabilities of mutant and non-mutant patients are equal, due to the stratified sampling for training. Because *P*_*learn*_(*X* = *mutant*) = 0.5 is greater than *P*_*indep*_(*X* = *mutant*) ≈ 0.1, additional scaling or stringency is required to avoid false positives from the learned classifier applied to the independent dataset.
2. Evaluating *P*_*indep*_(*X*|*I*_0_, *I*_1_, …, *I*_8_, *C*_1_, *C*_2_,…, *C*_11_) for each patient. Each of the eleven classifiers are evaluated on the patient’s central representative histology image and the octagon of eight surrounding images (Fig 4 panels B and C), to predict SPOP mutation. Because each classifier has been trained on only 10 mutants and 10 non-mutants, many classifiers will emit a biased prediction of 0 for all nine images or 1 for all nine images, lacking power to discriminate among the nine images. Empirically however, 1-3 classifiers tend to have greater variance in their predictions for each of the nine images of the patient. The classifiers with greater variance tend to correctly predict SPOP mutation state, presumably because the features learned from similar training data can distinguish image regions showing mutant image textures from regions showing non-mutant image textures. Such mutant and non-mutant regions should spatially cluster, i.e. a classifier should make the same SPOP mutation state prediction for adjacent images in a patient, otherwise the classifier is predicting noise. Moreover, classifiers with greater variance should agree which patches have high evidence of mutation and which patches have low evidence of mutation. Therefore, we take the classifier mean prediction, for all 11 classifiers in the ensemble, weighting more highly in this mean the predictions from classifiers with higher interpatch prediction variance. We define the weighted ensemble average as 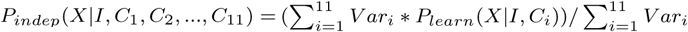 where we define 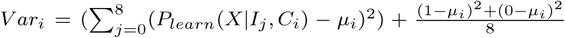 and 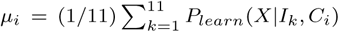, so that even clas sifiers that predict 0 for all images or 1 for all images still have some small weight. If, for instance, the mean predictions (*P*_*indep*_(*X*|*I*, *C*_1_, *C*_2_, …, *C*_11_)) suggest SPOP is mutated in each of 4 adjacent images, there is significant evidence for SPOP mutation at *α* = 0.05: of 9 images, 16 possibilities exist of 4 adjacent images being SPOP mutants and the other images being non-mutants, and 2^9^ = 512 mutant/non-mutant configurations possible, so probability of 4 adjacent images is 16/512 = 0.03125 < 0.05 = α. Thus for added stringency, one may take as the overall corrected *P*_*indep*_ the maximum of all adjacent 4-image cluster mean prediction minima: *P*_*indep*4_(*X*|*I*_0_, *I*_1_, …, *I*_8_, *C*_1_, *C*_2_, …, *C*_11_) = max(min_*i*∈*a*,*b*,*c*,*d*_(*P*_*indep*_(*X*|*I*_1_, *C*_1_, *C*_2_, …, *C*_11_))). However, due to the noisy nature of the mean predictions from classifiers trained on little data, a mutant 4-cluster may not occur, but additional adjacent images indicating mutation may offer supporting evidence. From the observation that a 4-cluster is two overlapping 3-clusters, we slightly loosen the correction stringency to *P*_*indep*3*pen*_(*X*|*I*_0_, *I*_1_, …, *I*_8_, *C*_1_, *C*_2_, …, *C*_11_) = *pen*(min(*P*_*indep*_(*X*|*I*_*a*_, *C*_1_, *C*_2_, …, *C*_11_), *P*_*indep*_(*X*|*I*_*b*_, *C*_1_, *C*_2_, …, *C*_11_), *P*_*indep*_(*X*|*I*_*c*_, *C*_1_, *C*_2_, …, *C*_11_))), ∀{*I*_*a*_, *I*_*b*_, *I*_*c*_} ∈ {*I*_0_, *I*_1_, …, *I*_8_} *s.t I*_*a*_ *adjacent I*_*b*_, *I*_*b*_ adjacent *I*_*c*_, where *pen*(…) returns the second-greatest value, or penultimate. In this way, *P*_*indep*3*pen*_ is identical to *P*_*indep*4_ for a 4-cluster, but additionally allows partially‐ or non-overlapping 3-clusters to suggest SPOP mutation. *P*_*indep*3*pen*_ is strictly greater than or equal to *P*_*indep*4_.

### S4. Implementation Details

We studied a TCGA cohort of 499 prostate adenocarcinoma patients, in particular the 177 patients that had both acceptable pathology images and SPOP mutation state available (Fig 1, Alg S1). SPOP mutation state was downloaded from cBioPortal [27]. After learning on TCGA data, we tested on an MSK-IMPACT dataset of 19 SPOP mutant patients and 119 non-mutant patients.

Microscope slides were scanned at 0.25 ± 0.003 microns per pixel *[μpp]*, using an Aperio AT2 scanner. The resulting SVS data file consists of multiple levels, where level 0 is not downsampled, level 1 is downsampled by a factor of 4, level 2 by a factor of 16, and level 3 by a factor of 32. From each level, 800×800px patches were extracted via the OpenSlide software library [30]. We refer to level 2 as low magnification and level 0 as high magnification. Level 2 approximately corresponds to a 10x eyepiece lens and 10x objective lens at the microscope when the scan is 0.5*μpp*. Our saliency predictor assumed 0.5*μpp* scans, though the scans here were 0.25*μpp*, but appeared robust.

Algorithm S1 describes data preprocessing for training. In prior work, we developed a patch saliency predictor [18]. A TMARKER classifier was trained to determine cell types [28]. We define the dominant tumor patch as having the maximum number of cancer cells of all 800×800px salient patches at low magnification. Within the 800×800px dominant tumor patch, we select an 800×800px patch at high magnification having the maximum number of abnormal cells. Malignant cells count 1 towards the maximum abnormal cell count, unknown cells count 0.5, and healthy cells count 0. The dominant tumor is explored in increments of 10 pixels, until the bounding box with the maximum number of abnormal cells is found. Whereas our saliency predictor operated on low-power tissue-level details of a patch, our SPOP predictor operates on high-power cell-level details of a patch. A patient is discarded from the study if the abnormal patch over-stained eosin blue, or is > 50% blurred, or is > 50% background, or is > 50% blood – as determined by visual inspection.

**Fig. S1.**
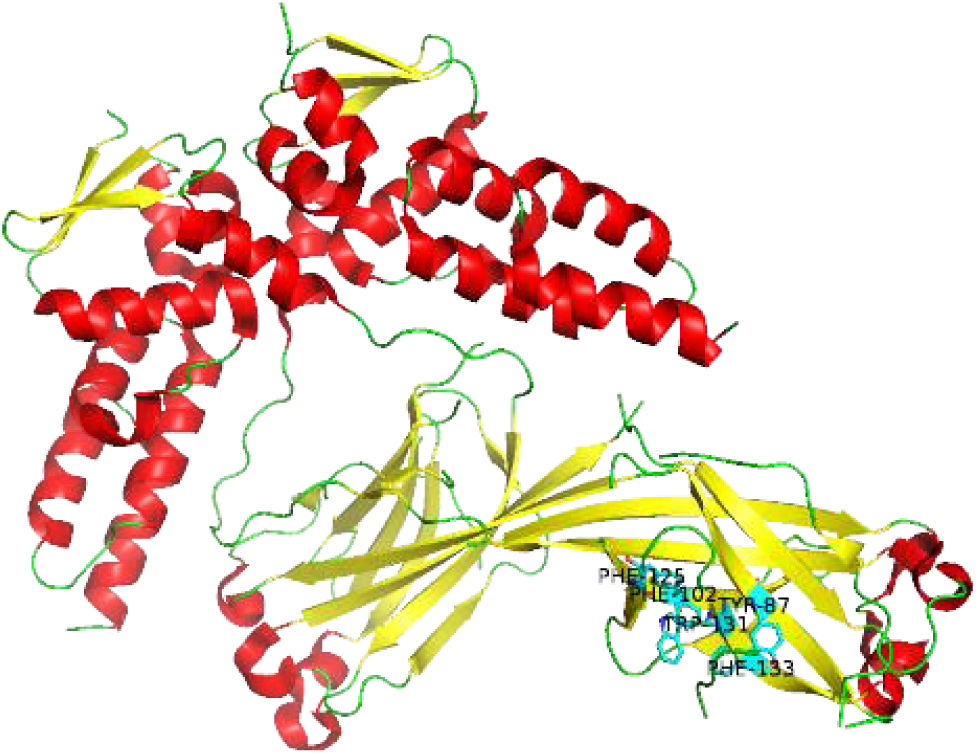
SPOP tertiary structure with mutated residues labeled, common to one site. Though most genetic aberrations affect the substrate-binding MATH domain in SPOP, our method’s performance appears robust to two of 20 patients having genetic changes outside this pocket. Mutation chr17:47677762 R368H in patient TCGA-EJ-7782-01 not shown and not in site. Deletion chr17:47699392 in patient TCGA-VP-A878 not shown and not in site. PDB structure 3HQI.

Algorithm S2 describes the neural network ensemble for training. This procedure allows learning to occur despite a mere 20 patients having an SPOP mutation, compared to 157 not having an SPOP mutation. Additionally, having 5 ensembles – each trial yielding one ensemble of 11 ResNets – allows a CI in predicted SPOP mutation state to be calculated for both the generalization error estimate during training and any future patient (Algorithm S3). Moreover, such a large number of ResNets can fully sample the SPOP nonmutant patients, while each ResNet is still trained with an equal proportion of SPOP mutants and non-mutants. We use two neural network architectures, both the published ResNet-50 architecture (*r*50 in Alg S2) and our custom ResNet-50 with a 50% dropout [4] layer with an additional 1024 fully-connected layer as the top layer (*drp*5 in S2, and shown as Drop50 in Fig 4). In practice, at least one architecture tended to have validation accuracy ≥ 0.6. Architecture diversity may increase intra-ensemble ResNet variance, and the decorrelation in errors should average out in the ensemble. Trials 0, 2, and 3 used two Drop50 learners and nine ResNet-50 classifiers (Fig 4, Table S1). Trial 1 had six and five, respectively. Trial 4 had three and eight. Trials 1 and 4 had the worst performance by AUROC (Fig 5), both had at least one ResNet-50 predictor with 0.3 or worse test set accuracy (Table S1)), and both had more than two Drop50 learners in the final ensemble for the trial. For challenging draws of training, validation, and test sets, Drop50 learners may slightly outperform ResNet-50 learners, though generalization accuracy may remain low due to the clustering of the data in the draws and limited sample sizes.

ResNets (both ResNet-50 and Drop50 architectures) were trained on 10 mutant and 10 non-mutant patients, so with data augmentation each ResNet was trained on 21600 images total. For each training set, a validation set of 5 mutant and 5 non-mutant patients was used and did not overlap with the training set, so with data augmentation each ResNet was validated against 10800 images. The test set for an ensemble consistent of 5 mutant and 5 non-mutant patients which were not augmented and did not overlap with any training or validation set for any classifier in the ensemble, so each ensemble was tested against 10 images.

There is remarkable variability in test accuracy among the testable models from each Monte Carlo cross validation run (Table S1). If the test set is drawn from approximately the same distribution as the validation set, where ResNet validation accuracy is 0.6+ and ResNets are uncorrelated, then we can expect 6 of the 11 ResNets (6/11 < 0.6) in an ensemble to correctly predict SPOP mutation state on average. In this way the ensemble is a strong learner based on ResNet weak learners [19, 20]. Through ensemble averaging, the mean SPOP mutation probability is computed over all 11 constituent ResNets, to provide the final probability from the ensemble.

**Fig. S2.**
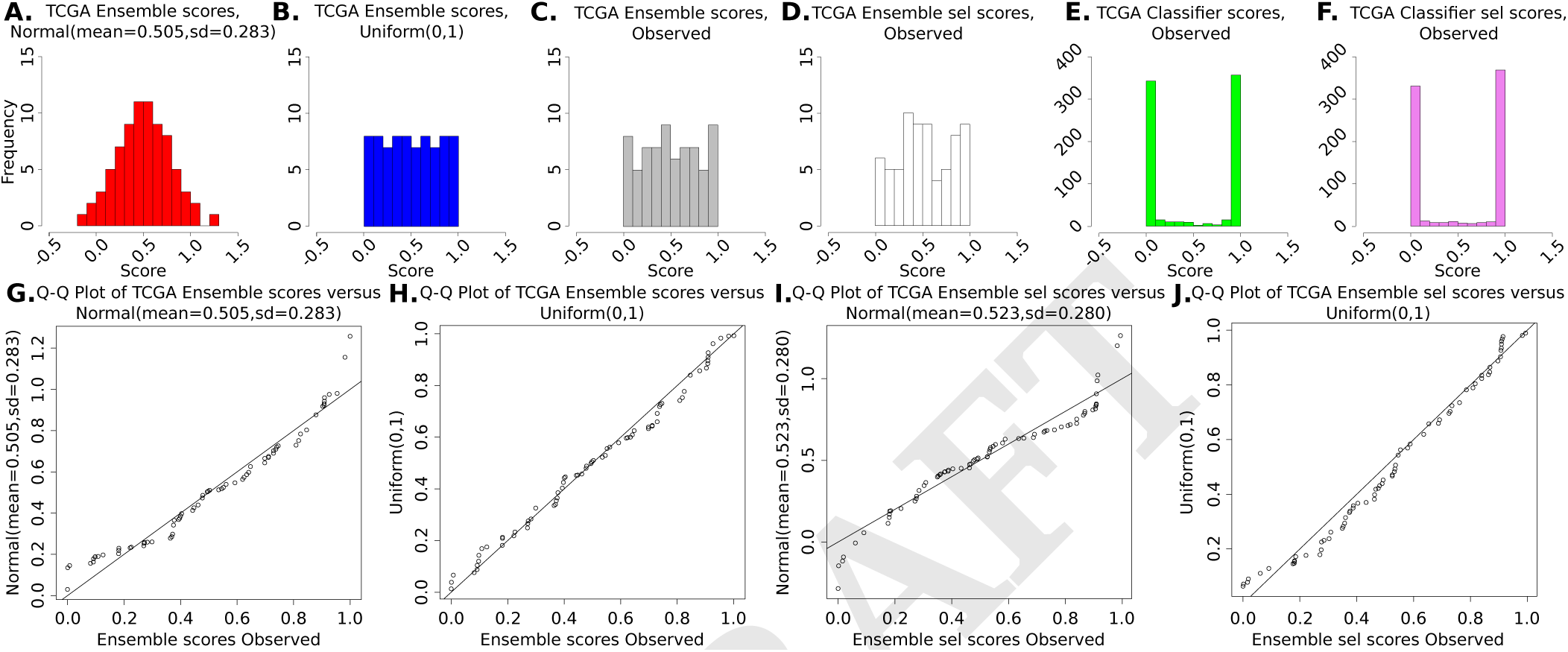
TCGA classifier and ensemble mutation prediction scores, showing ensemble scores are uniform random variables representing the mutation probability when the mutant and non-mutant classes are equally sampled, despite the sharply bimodal score distribution of individual classifiers (Table S1). *Compared to the bimodal distribution of single classifiers, this uniform random distribution enables valid confidence interval calculation over ensemble mutation prediction scores* and additionally does not suffer confounds such as p-value inflation, e.g. a disproportionate number of ensemble mutation prediction scores being close to zero. **Panel A:** Normal distribution of scores with mean and stdev of TCGA-trained ensemble prediction scores, showing the distribution domain extends beyond the valid [0,1] domain of mutation prediction scores. **Panel B:** Uniform distribution of scores, with all scores in the valid [0,1] domain and none outside. **Panel C:** TCGA-trained ensemble scores, following Uniform distribution (KS Test p=0.9792). Corresponds to Fig 4 panel A, second from right. **Panel D:** TCGA-trained tuned ensemble scores, following a Uniform distribution (KS Test p=0.4687). Corresponds to Fig 4 panel A, far right. The excess of predictions around 0.5 is an artifact of the tuning process, which tends to select classifiers for the ensemble such that the ensemble prediction is at least 0.5 for mutants and less than 0.5 for non-mutants if the ensemble prediction was incorrect before tuning. See also Fig S4 for tuned ensemble performance. **Panel E:** TCGA-trained classifier scores, following sharply bimodal distribution of 0 (non-mutant) or 1 (mutant) predictions. **Panel F:** TCGA-trained classifiers selected for the tuned ensemble are also bimodal. **Panel G:** Q-Q plot showing outliers, indicating TCGA-trained ensemble scores do closely follow a Normal distribution. **Panel H:** Q-Q plot showing close linear relationship, indicating TCGA-trained ensemble scores follow a Uniform distribution. **Panel I:** Q-Q plot showing outliers, indicating TCGA-trained tuned ensemble scores do not closely follow a Normal distribution. **Panel J:** Q-Q plot showing close linear relationship, indicating TCGA-trained tuned ensemble scores follow a Uniform distribution.

**Fig. S3.**
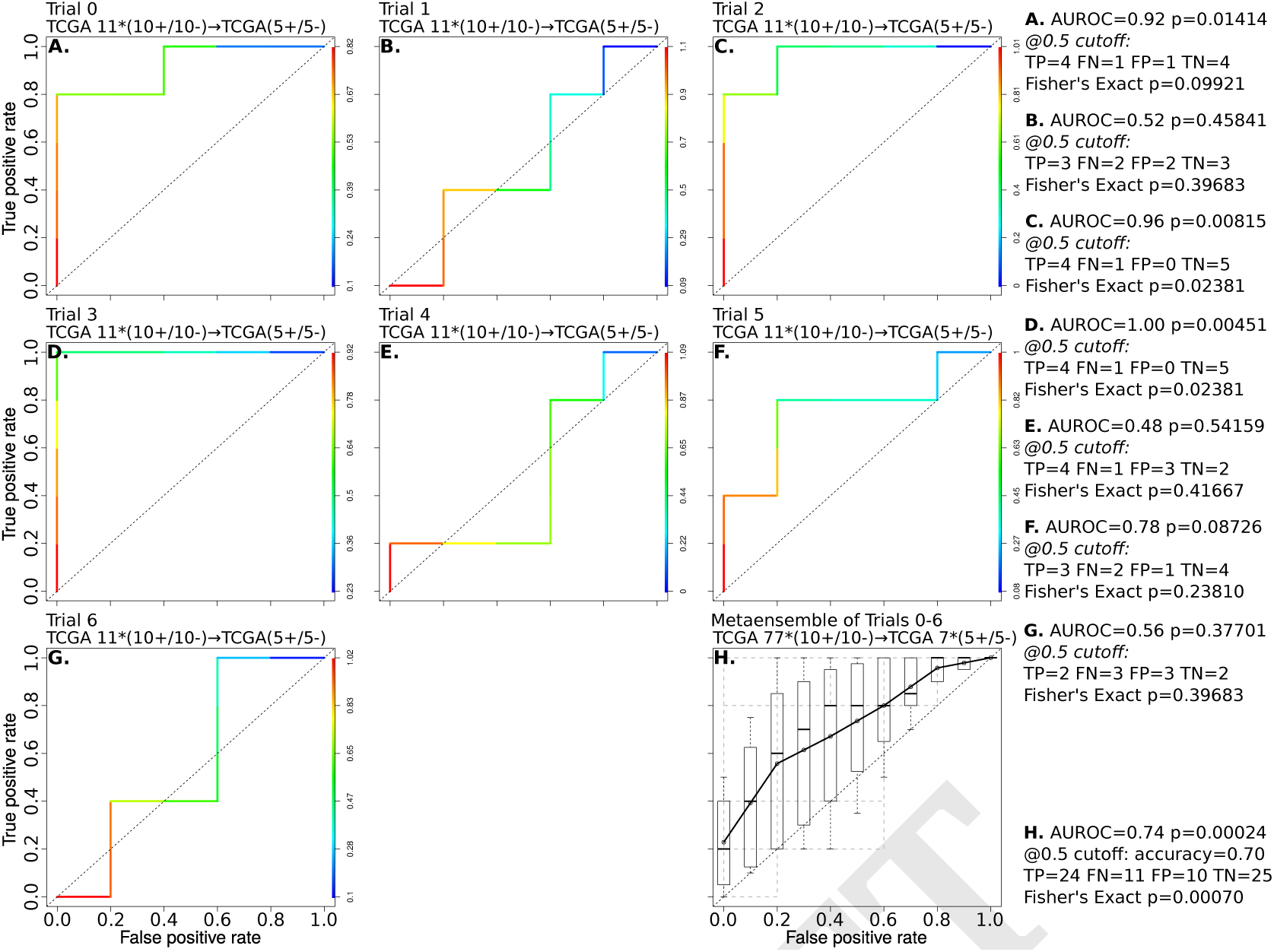
Ensemble performance, for comparison to ensemble performance in Fig 5 panel C1. Classifiers in ensemble selected by highest validation accuracy. This estimates performance on unseen data. Performance for the classifiers instead selected by highest test set accuracy is shown in Fig S4.

**Fig. S4.**
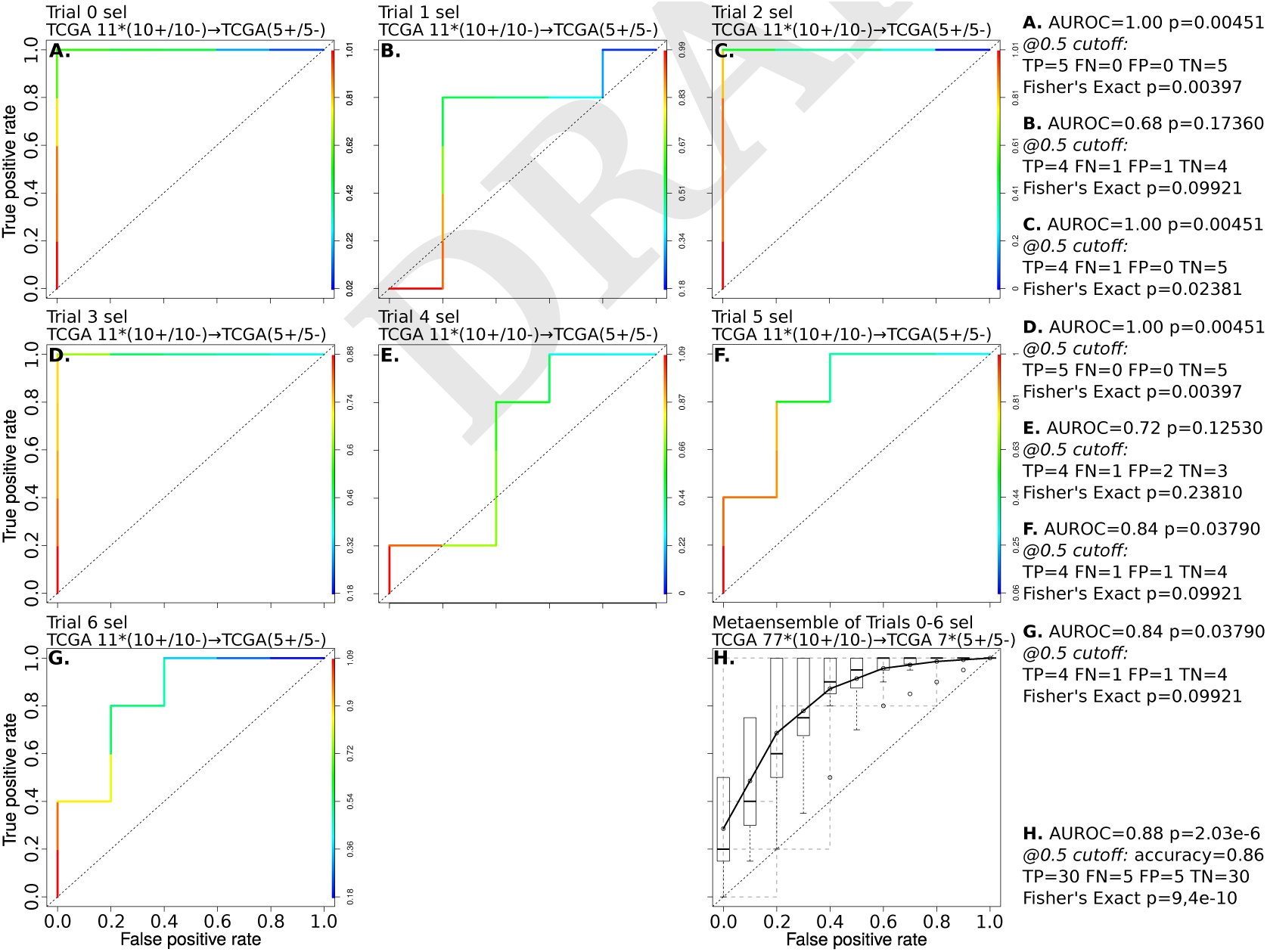
Tuned ensemble performance, for comparison to ensemble performance in Fig 5 panel C. The tuned ensembles are used for prediction on the independent MSK-IMPACT cohort (Fig 5 panels C2 and C3). Tuning provides limited additional training for each ensemble via model selection on the corresponding TCGA test set. Tuned ensemble mutation prediction distributions shown in Fig S2.

**Table S1.**
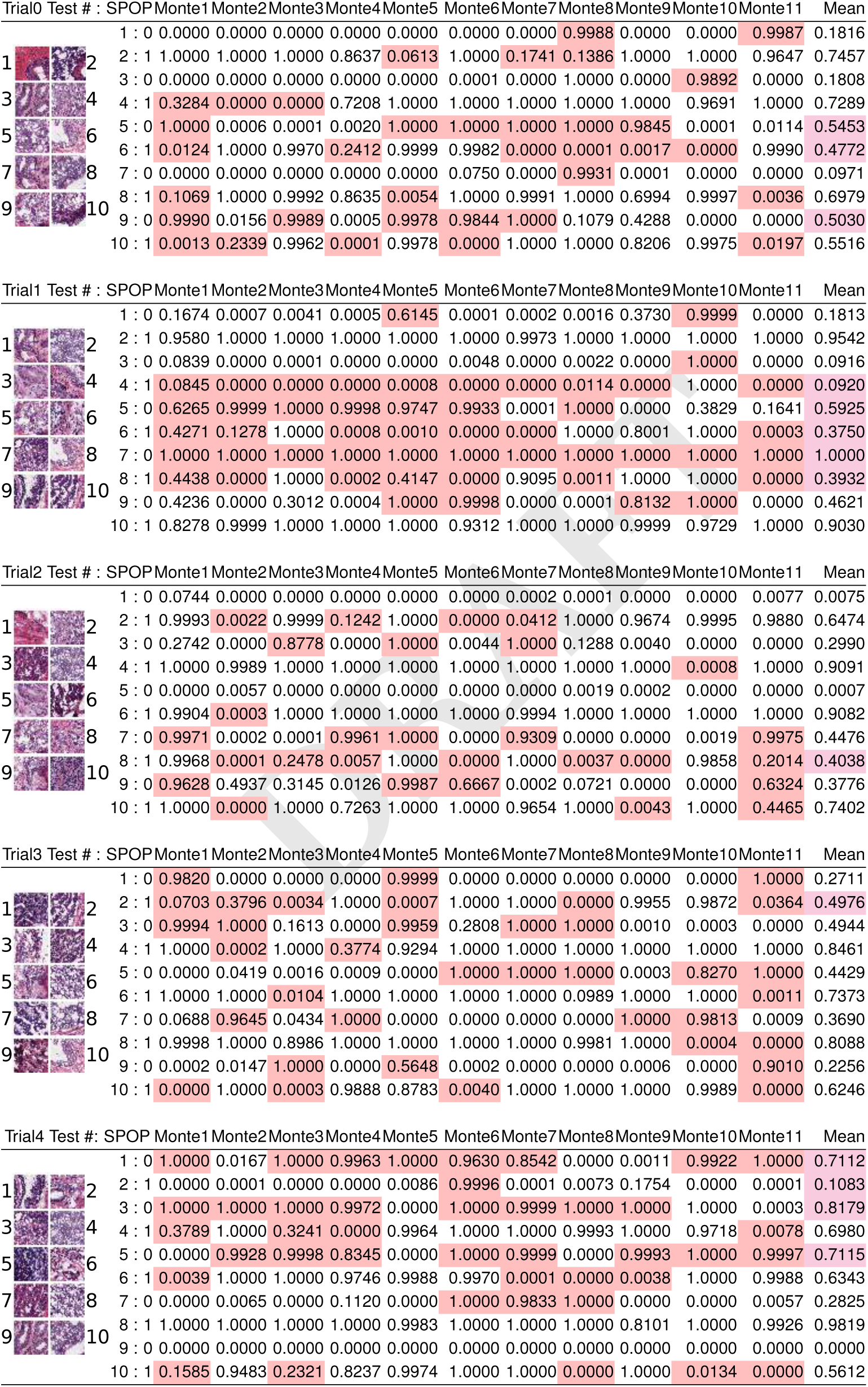
TCGA test set images and accuracies, with prediction errors from single residual networks highlighted in red and prediction errors from residual network ensembles highlighted in magenta. New 20-patient training and 10-patient validation sets are drawn for each Monte Carlo cross validation run against the trial’s 10-patient test set. The far right column is used to calculate the mean classifier prediction, which is used as the trial’s ensemble prediction. The Receiver Operating Characteristic of these ensembles in shown in Fig S3. The seven ensemble predictions are used to calculate a confidence interval of generalization accuracy, shown in Fig 4 panel A second from right in gray and red. Individual classifier predictions follow a sharply bimodal distribution of 0 (non-mutant) or 1 (mutant), while the ensemble predictions follow a uniform random distribution (Fig S2). Table S1 is continued in Table S2, showing the final two enembles in the seven ensemble metaensemble trained on TCGA data.

**Table S2.**
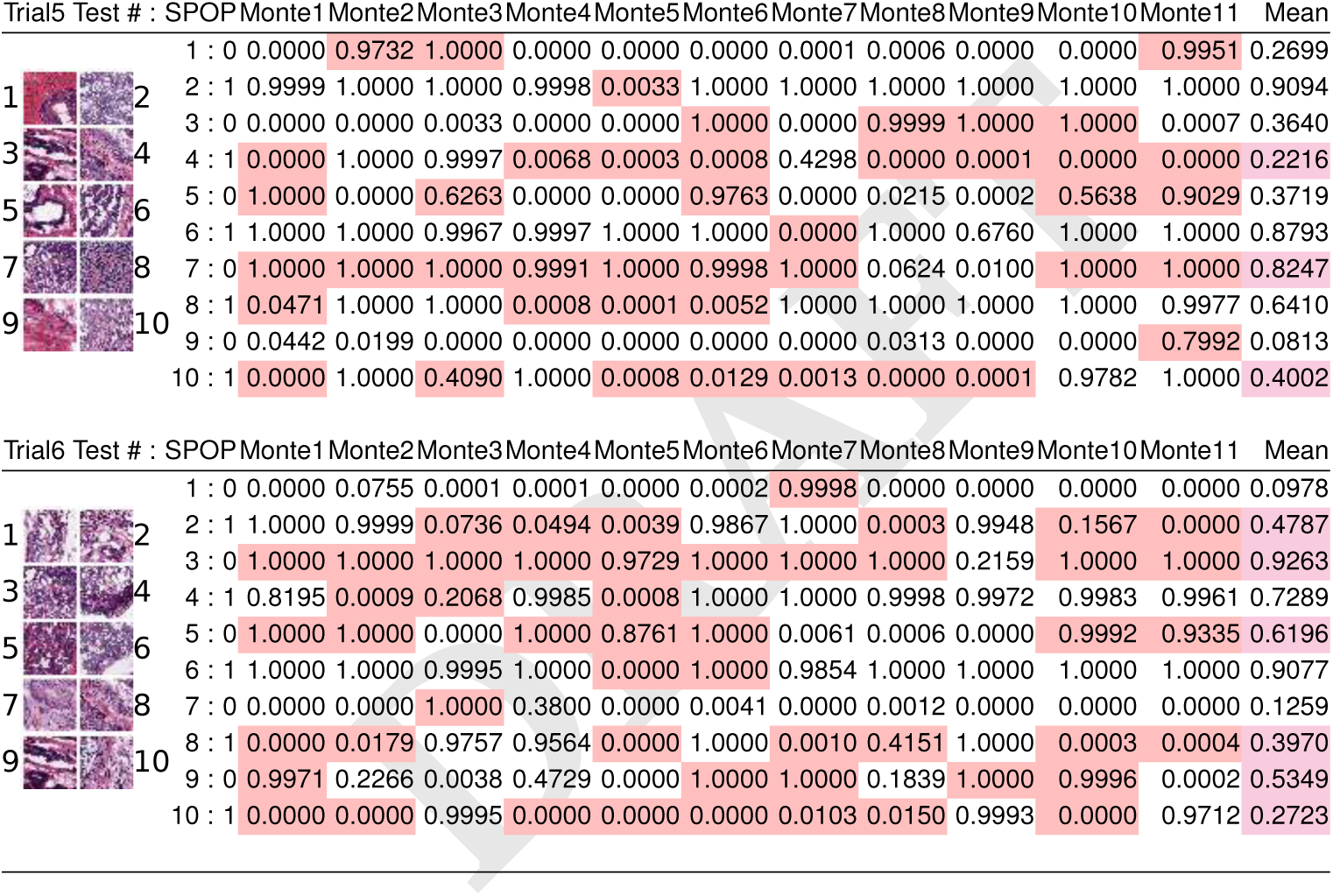
Continuation of Table S1, for trials 5 and 6, the final two ensembles in the TCGA-trained metaensemble.

## Footnotes

1 TCGA data courtesy the TCGA Research Network http://cancergenome.nih.gov/

2 We used the R boot library for basic bootstrap CIs. Canty and Ripley 2016: https://cran.r-project.org/web/packages/boot/index.html

3 We used Kost’s Method from R package EmpiricalBrownsMethod https://www.bioconductor.org/packages/devel/bioc/vignettes/EmpiricalBrownsMethod/inst/doc/ebmVignette.html

